# Adaptive evolution of *Candida maltosa* improves the bioconversion of depolymerized plastic feedstock by targeting biosurfactant production

**DOI:** 10.1101/2025.10.11.681799

**Authors:** Efrain Rodriguez-Ocasio, Kimia Noroozi, Ammara Khalid, Jessica Brown, Robert C. Brown, Mark A. Blenner, Laura R. Jarboe

## Abstract

Thermal oxo-degradation (TOD) of plastic can transform plastic waste into fermentable feedstocks. Bioconversion of the TOD products could be used as feedstocks in a biorefinery concept and lead to new avenues for plastic waste upcycling. Previous work demonstrated this concept with high density polyethylene (HDPE), the most abundant type of plastic, and identified the nonconventional yeast *Candida maltosa* as a promising candidate for this application. Here we describe the evolution of an improved strain of *C. maltosa* and characterize the uptake mechanisms of TOD products from HDPE (TOD_HDPE). Batch cultures in series passaged at the mid-exponential growth phase applied a selective pressure for faster growth and resulted in a >100% increase in specific growth rate when using TOD_HDPE as a carbon source. The evolved strain was compared to the parent strain to identify the cellular and biochemical changes associated with the improved phenotype and the uptake mechanisms involved in the bioconversion of TOD_HDPE. This comparison found that *C. maltosa* secretes biosurfactants capable of solubilizing hydrocarbons. The adaptive evolution resulted in changes in biosurfactant production that translated to improved emulsification of alkanes and increased solubilization of fatty alcohols and alkanes. In addition to the changes in metabolites, the study identified increases in membrane permeability associated with a reduction in ergosterol that may also play a role in the improved phenotype. These findings support the development of *C. maltosa* and other potential microbial cell factories for plastic biorefineries and may inform future design strategies.

## Introduction

Global plastic production is rapidly increasing, but existing waste management strategies have proven inadequate and most of the plastic waste ends up in landfills and the environment [1]. Plastic pollution causes physical, chemical, and biological threats to organisms and human health [2]. There is a pressing need for new technologies to prevent plastic accumulation in the environment and microbial conversion into value-added products could be an impactful technology in the fight against pollution. Biodegradation is not fast enough for the industrial processing of most plastics [3], [4]. In contrast, thermal oxo-degradation rapidly depolymerizes and oxidizes the polymers into fermentable products, making it possible for microorganisms to metabolize the carbon from plastic wastes [5]. This approach was previously demonstrated with polyethylene as the plastic waste and the non-conventional yeast *Candida maltosa* as the microbe [5].

The products from thermal oxo-degradation of high-density polyethylene (TOD_HDPE) are a waxy mixture of mostly solid hydrophobic molecules, which poses a challenge for fermentation because low solubility limits the availability of the carbon source in the aqueous media. However, *C. maltosa* still showed substantial growth even in the absence of emulsification or alternative bioprocesses like solid-state fermentation [5]. Thus, *C. maltosa* has the potential to become a biomanufacturing platform for hydrophobic feedstock utilization. Our main interest is in repurposing the biorefinery concept for plastic waste upcycling, but this organism could also be useful for bioremediation and treatment of fats, oils and greases (FOGs) [6].

The substrate range for *C. maltosa* is mid-to long-chain hydrophobic hydrocarbons and lipids [7]. However, the uptake mechanism of hydrophobic substrates that present mass transfer limitations is poorly studied in this and other organisms. Current understanding on this topic is based on investigations of the oleaginous yeast *Yarrowia lipolytica* [8], [9]. However, *C. maltosa* shows superior growth in thermal oxo-degradation products of polyethylene and the two species show differences in response to hydrophobic substrates [5]. For example, *C. maltosa* forms canals in the form of proteinaceous supramolecular structures across the cells wall in response to alkanes while *Y. lipolytica* does not [10], [11]. Understanding how *C. maltosa* overcomes the mass transfer limitations and utilizes the out-of-phase mixed feedstock as its sole carbon source can inform future strain engineering and bioreactor design.

In this study, we used adaptive laboratory evolution to improve the growth rate of *C. maltosa* on thermally oxo-degraded polyethylene. Adaptive laboratory evolution is a powerful tool within the Design-Build-Test-Learn framework of metabolic engineering [12], [13]. The application of a selective pressure allows isolation of cells with physiological adaptations that enable faster growth than those without such adaptations. Comparison of the evolved strain and the wild-type parent strain can identify changes in substrate range, morphology, subcellular structures, membrane properties and biosurfactant production. Specifically, our findings paint a picture of the mechanisms involved in the uptake and metabolism of thermal oxo-degradation products that can possibly be extrapolated to other hydrophobic feedstocks.

## Materials and Methods

### Thermally Oxo-degradation of Plastics

Wax from thermally oxo-degraded high-density polyethylene (TOD_HDPE) was obtained as previously described using an electrically heated, semi-batch tubular reactor (Figure 1), as described in Reference [5]. The reactor is equipped with a feed auger that continuously drops feedstock into a heated 2.54 cm stainless steel tee, where devolatilization occurs. Prior to the reactor tee is a gas pre-heat zone capable of heating the incoming mixture of nitrogen and air controlled by individual mass flow controllers. Following the devolatilization zone is a 38 cm long, 3.5 cm diameter tubular reactor, named the “gas phase oxidation zone”, in which vapors undergo further cracking and oxidation. Passive cooling and an externally heated in-line 40 µm impact filter were utilized for collection of wax using minimal energy. Product was collected in a sterile pre-weighed bottle affixed to the collection system. Vapors not condensable using this configuration were vented.

**Figure 1.**
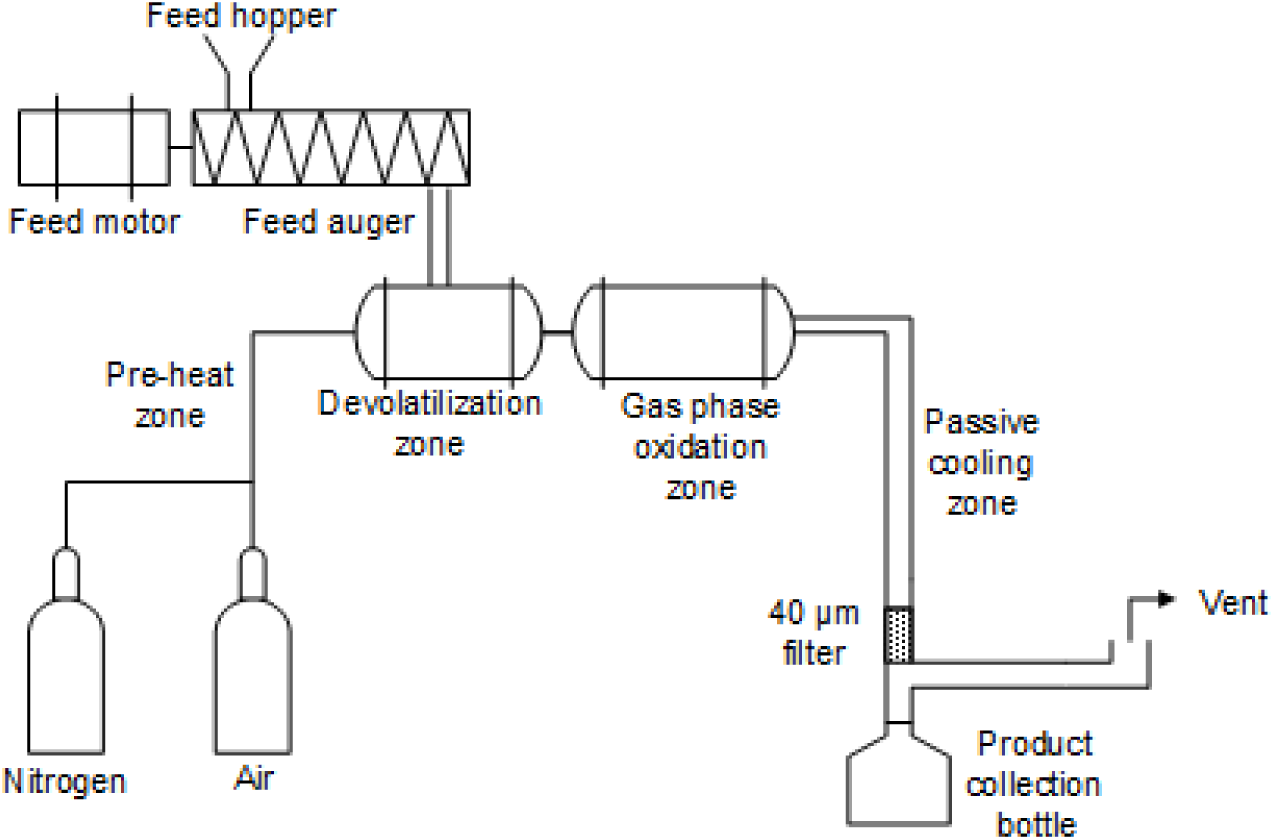
Electrically heated, continuous thermal reactor used to perform thermal oxo-degradation of plastic. Heat tapes surrounding the reactor zones are not illustrated. [5]

Here, the feed hopper was loaded with 450 g of injection-grade HDPE pellets (Advanced Production Systems, Ohio, US). Trials were performed using 100% w/w HDPE and 50:50 HDPE:PP as feedstock. Using external heaters, the pre-heat zone was maintained at 525°C, the devolatilization zone at 500°C, and the gas-phase oxidation zone at 425°C. Nitrogen and air flow were initiated at 2.25 standard liters per minute (SLPM) and 0.75 SLPM, respectively. The heated vapor residence time was estimated as 3.8 seconds based on the gas flow rate and the internal volume of the combined devolatilization and gas-phase oxidation zone of the reactor. Plastic pellets were fed at a rate of 100 g/h and 3.5% of the air required for the complete combustion of the feed was metered into the system. After all plastic pellets were fed, the system was allowed to cool, and the bottle containing product was weighed. Yield of product was 67 % w/w on a fed-plastic basis.

### TOD Product Characterization

Fourier-transform infrared spectroscopy (FTIR) was used to identify the functional group composition of waxes and spent media components and to calculate the carbonyl index of waxes [14]. Using a Thermo Scientific Smart iTR Nicolet iS10 instrument fitted with a diamond window, samples were scanned 64 times from 400 cm^-1^ to 525 cm^-1^ with resolution of 4 cm^-1^. The baseline was normalized after the spectra was produced.

The molecular weight distribution of the wax was determined by using gel permeation chromatography (GPC). Approximately 25 mg of wax was dissolved in 10 mL of tetrahydrofuran (THF) and filtered through a 0.45 μm PTFE syringe filter into 2 mL vials. Analysis was performed using a Dionex Ultimate 3000 high performance liquid chromatograph fitted with one MesoPore column (3 μm inner diameter, 300 x 7.5 mm; 200-25,000 Da) and two Agilent PLgel columns (3 μm inner diameter, 300 x 7.5 mm; 100-4000 Da) in series. THF was used as the mobile phase at a flow rate of 1.0 mL/min at 25 °C. A Shodex Refractive Index (RI) and a diode array detector (DAD) were used for measurements. Polystyrene standards were used for instrument calibration.

### Adaptive Laboratory Evolution

*C. maltosa* NRRL Y-17677 was obtained from the USDA ARS Culture Collection, rehydrated in YPAD media (10 g/l yeast extract, 20 g/l peptone, 0.4 g/l adenine hemisulfate, and 20 g/l dextrose, pH 5.5) and stored in a 25 % v/v glycerol suspension at -80oC. A single colony was selected from a YPAD agar plate streak-out and incubated in 3 mL BD Difco YNB minimal media without amino acids at pH 5.5 and supplemented with 2.0% w/v glucose (described henceforth as YNB + 2% glucose) at 30oC and 250 rpm for 24 hours. The preculture was washed by pelleting the cells by centrifugation at 4415 × *g* for 5 minutes and resuspending in the same volume of YNB without any carbon source. The directed evolution was carried out simultaneously in three replicates. Each replicate started in a 250 mL baffled flask with 50 mL of YNB without amino acids supplemented with 0.5% w/v TOD_HDPE (YNB + 0.5% w/v TOD_HDPE). The TOD_HDPE remained as a solid floating in the media with a consistency similar to butter. Each flask was inoculated with the washed preculture at a starting OD_600_ of 0.1. The cultures were incubated at 30oC with 250 rpm agitation in a MaxQ 6000 incubated shaker. When a culture reached an OD_600_ of 0.6, a 100 μL aliquot was transferred to a new flask with fresh 50 mL YNB + 0.5% w/v TOD_HDPE. The OD_600_ measurements were blanked against TOD_HPDE no cell controls to account for the contribution of TOD_HDPE particulate. The serial cultures were performed for 14 batches over 427 hours.

### Evolved Strain Isolation

The evolved population was streaked out on a YPAD plate and incubated at 30oC for 48 hours. After incubation, 20 distinct colonies were selected as candidate strains and individually cultured in YNB + 2% w/v glucose. For the first phase of selection, the 20 candidates were co-cultured in groups of five in YNB + 0.5 w/v TOD_HDPE at 30oC and 250 rpm agitation. The specific growth rate (μ) from each co-culture was determined using the following formula:

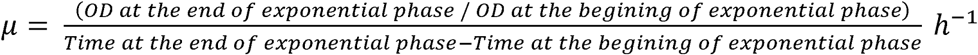

In the second phase of selection, the co-culture with the highest specific growth rate was selected and the specific growth rate of the individual candidates was determined using the same methodology. The strain with the highest growth rate from the five candidates was selected as the final evolved strain.

### Strain Comparison in Model Compounds

The growth profile of the evolved strain of *C. maltosa* was studied using the model compounds and methodology from a previous survey of non-conventional yeast metabolism that used the parent strain [7]. Briefly, 14 mL culture tubes with 3 mL of YNB without amino acids and 0.050 M of a focal model compound were prepared in triplicates for 18 model compounds and for both the parent and evolved strains. One set of model compound cultures was inoculated with the parent strain and the other with the evolved strain, both with a starting OD_600_ of 0.1. All conditions were incubated at 30oC with 250 rpm agitation for 40 hours.

### Strain Comparison in Alternative Carbon and Nitrogen Sources

Strains were grown in TOD_HDPE and TOD products from a 1:1 HDPE and polypropylene mixture (TOD_HDPE/PP) as sole carbon sources were prepared in 250 mL baffled flask with 50 mL of YNB without amino acids supplemented with 0.5% w/v of the corresponding feedstock. Urine was tested as an alternative nitrogen source to the ammonium sulfate (AmSO_4_) in YNB. Strains were prepared in 250 mL baffled flask with 50 mL of YNB without amino acids and without AmSO_4_, supplemented with 15% v/v urine (Lee Biosolutions, 991-03-P-FNF) and 2% w/v glucose. Each flask was inoculated with a single strain at a starting OD_600_ of 0.1 and incubated at 30oC with 250 rpm agitation. Growth was monitored by measuring the OD_600_.

### Model Compound Solubility

The parent and evolved strains of *C. maltosa* were cultivated in triplicates in YNB without amino acids supplemented with 2.0% w/v glucose for 48 hours. No cell controls with 2.0% w/v glucose TOD_HDPEwere also incubated for 48 hours in triplicates. After incubation, all conditions were filtered using a 0.2 µm filter. Three 600 μL aliquots for each biological replicate were collected, weighed and stored to serve as baselines. Subsequently, 1-tetradecanol, 1-octadecanol, n-tetradecane and n-octadecane were added individually to aliquots of each type of spent media at 0.5% w/v, followed by vortexing for 10 minutes and filtration. Each substrate/media combination was subjected to GC-MS analysis in triplicate.

A standard solution of 1.0 mg/mL 1-tridecanol in 3:1 v/v mixture of butanol and hexane (hereafter referred to as solvent) and spiked into each sample as an internal control at a final concentration of 0.08 mM. Extractions for GC-MS were conducted by mixing with equal volume of solvent, vortexing for 10 minutes, centrifugation at 15,000 x g for 10 minutes and transferring the organic layer to GC vials. The samples were dried in a speed-vac concentrator for 10 hours and then trimethylsilylation was performed by adding 100 μL of bis-trimethyl silyl trifluoroacetamide with 1 % v/v Trimethylchlorosilane for 30 min at 60 °C [15].

Derivatized samples were analyzed by GC-MS at the Iowa State University W.M. Keck Metabolomics Research Laboratory (RRID:SCR_017911). GC-MS analyses were performed with an Agilent 6890 gas chromatograph coupled to a model 5973 Mass Selective Detector (Agilent Technologies, Santa Clara, CA). The column used was HP-5MS 5% Phenyl Methyl Silox with 30 m × 250 µM × 0.25 µm film thickness (Agilent Technologies). One microliter of sample was injected with the inlet operating in splitless mode and held at a constant temperature of 280 °C. The oven temperature was programmed as follows: an initial temperature of 70 °C was increased to 250 °C at 15 °C/min and before being further increased at 20 °C/min to a final temperature of to 320 °C which was held for 8.5 minutes. Helium was used as a carrier gas at a flow rate of 1 mL/min. The MS transfer line was held at 280 °C. Mass Spectrometry detection was performed using electron ionization at 70 eV and source temperature and quadrupole temperature were set at 230 °C and 150 °C, respectively. The mass data were collected in the range from m/z 40 to m/z 800.

Identification and quantification were conducted using AMDIS (Automated Mass spectral Deconvolution and Identification System, National Institute of Standards and Technology (Gaithersburg, MD) with a manually curated retention indexed GC-MS library with additional identification performed using the NIST17 and Wiley 11 GC-MS spectral library (Agilent Technologies, Santa Clara, CA). Final quantification was calculated by integrating the corresponding peak areas relative to the area of the internal standard. Raw data were normalized to the mass of samples used.

### Surface Tension and Emulsification Index

Cells were cultured in triplicates in YNB + 0.5% w/v TOD_HDPE or 2% w/v glucose in 1 L baffled flasks with 200 mL working volume at 30oC with 250 rpm orbital agitation for 148 and 48 hours respectively. No cell controls were also incubated in triplicates for both the glucose and TOD_HDPE media. After incubation, the cultures were centrifuged at 3500 x g for 10 minutes and the supernatants were filtered through a 0.2 µm cellulose nitrate membrane. The surface tension of the filtered spent media was measured using the KIMBLE Surface Tension apparatus (DWK Life Sciences, Germany), based on the height of the liquid in a capillary tube. The emulsification index for each condition was measured using a modified version of the methods described in References [16], [17]. Iqbal et al., 1995 and Zarur Coelho et al., 2010Specifically, the filtered spent media was mixed at 1:1 volume ratio with the specified solvent, vortex-mixed for 10 minutes and allowed to rest at room temperature for 1 hour. The emulsification index was estimated as the height of the emulsion formed divided by the total height of the two-phase liquid.

### Biochemical Characterization of Biosurfactants

The saponification test was performed in triplicates as previously described [18]. Specifically, a 0.5 mL sample of filtered spent media was mixed with 1 mL of 20 % w/v NaOH and 0.5 mL of 100 % ethanol and placed in a boiling water bath for 15 minutes, followed by the addition of 5 mL of deionized H_2_O and then vigorous shaking. Samples were then visually inspected to note the presence of absence of froth, indicating negative or positive results, respectively. Positive controls were prepared with a drop of olive oil in 0.5 mL media and negative controls with only media.

Carbohydrates were detected using the anthrone test [19]. Triplicate 1 mL samples of filtered spent media were mixed with 5 mL of 2 mg/mL anthrone solution and incubated at 90 °C for 17 minutes. The samples were cooled to room temperature and the absorbance at 620 nm was measured. The presence of blue/green color with an absorbance significantly higher than that of a negative control (no cell control media) indicates a positive result. Positive controls were prepared with glucose.

The rhamnose test was performed to detect rhamnolipids [20]. Diethyl ether extracts of the spent media were dried under nitrogen and dissolved in ultrapure water. A 100 μL aliquot of the resuspension was mixed with 900 μL of 0.19 % w/v orcinol (Thermo Scientific Chemicals) in 53% v/v sulfuric acid and incubated at 80 °C for 30 minutes. After cooling to room temperature, the absorbance was measured at 421 nm and compared to a standard curve.

### Proteomics

Strains of *C. maltosa* were cultivated in two sets of triplicates in TOD_HDPE as described above. The cells from one set of triplicates were harvested during the exponential growth phase and the second set was allowed to reach stationary phase before filtration through a 0.2 µm membrane. The filtered spent media and yeast pellets were harvested and treated separately. The spent media samples were transferred to 1 kDa dialysis tubes (Cytiva 80648394) and placed in a 40 X volume deionized water reservoir with magnetic stirring at 4 °C overnight for analysis of secreted proteins. The yeast pellets were washed three times with cold phosphate-buffered saline (PBS) at pH 7.4 before resuspending approximately 150 mg in 4 mL of EZLys™ Yeast Protein Extraction Reagent (BioVision) and adding 10 µL of 1X EZBlock™ Protease Inhibitor Cocktail. The samples were incubated for 30 minutes at 30oC and 250 rpm agitation, then vortex for 30 seconds and centrifugation at 12,000 x g at 4oC for 15 minutes. The supernatants were collected and dialyzed in the same manner as the filtered spent media.

The buffer-free protein extracts were analyzed at the Iowa State University Protein Facility. The crude protein extracts were digested in solution with trypsin/Lys-C at a 1:25 enzyme:protein ratio. After digestion, peptide retention time calibration standard (Pierce part#88320) was added as an internal control. The peptides were separated by liquid chromatography and analyzed by MS/MS using the Q Exactive™ Hybrid Quadrupole-Orbitrap Mass Spectrometer. Raw data was analyzed with Thermo Scientific’s Proteome Discoverer Software and using Mascot to search against public databases for identification. The *Candida maltosa* Xu316 genome was used as reference. The relative abundance for each protein was normalized to internal standards.

### Membrane Permeability

Cells were grown to stationary phase in glucose and in TOD_HDPE individually, using the same methodology previously described above. Membrane permeability was measured in triplicates using a modified version of the membrane permeability fluorescence assay from [21]. Cells were centrifuged at 4415 x g for 5 minutes and the pellets were resuspended in 0.1 M Tris-HCL buffer pH 7.5 and normalized to an OD_600_ of 0.5. SYTOX Green (Invitrogen S7020) was added to each sample at a final concentration of 10 µM. A 200 µL aliquot from each condition was loaded into a flat-dark bottom 96-well plate. Bulk fluorescence was quantified in a SynergyHT BioTek microplate reader utilizing a filter of 485/20 nm and 516/20 nm for excitation and emission wavelengths. Fluorescence was monitored for 20 minutes until the measurements stabilized. The steady-state bulk fluorescence values were normalized by subtracting the bulk fluorescence of cells without the SYTOX dye and dividing by the OD_600_ of each condition.

### Microscopy

Cells were preserved with fixative for 24 hours with 1% w/v paraformaldehyde, 3% w/v glutaraldehyde in 0.1 M sodium cacodylate buffer, pH 7.2 at 4 °C to immobilize cellular structure in life-like state. The cells were then subjected to three 10-minute washes in cacodylate buffer and post fixed with 2% w/v potassium permanganate for 1 hour at room temperature. Subsequently, the cells were washed with deionized water 3 times for 15 minutes each, and en bloc stained using 2% w/v uranyl acetate in distilled water for 1 hour. Cells were washed in distilled water for 10 minutes and dehydrated through a graded ethanol series (25, 50, 70, 85, 95, 100% v/v) for 1 hour each step then further dehydrated with three changes of pure acetone, 15 minutes each, and infiltrated with Spurr’s formula epoxy resin (Electron Microscopy Sciences, Hatfield PA) with graded ratios of resin to acetone until fully infiltrated with pure epoxy resin (3:1, 1:1, 1:3, pure) for 6-12 hours per step. Cells were placed into capsules and were polymerized at 70 °C for 48 hours. Thick sections (1.5 µm) were made using a Leica UC6 ultramicrotome (Leica Microsystems, Buffalo Grove, IL) and stained with EMS Epoxy stain (a blend of toluidine blue-O and basic fuchsin). Thin sections were made at 50 nm and collected onto single slot carbon film grids. SEM and TEM was carried out by the Roy J. Carver High Resolution Microscopy Facility at Iowa State University using a 200 kV JEOL JSM 2100 scanning transmission electron microscope (Japan Electron Optics Laboratories, USA, Peabody, MA) with a GATAN One View 4K camera (Gatan inc., Pleasanton, CA).

### Metabolomics

Cells were grown in triplicates TOD_HDPE for 168 hours at 30oC and 250 rpm. The cells were pelleted by centrifugation and the supernatant was filtered with a 0.2 µm membrane. The cells were washed by resuspending in PBS pH 7.0, pelleting, discarding the supernatant and repeating the steps two more times. A non-targeted metabolite analysis was conducted on the cells using GC-MS and on the filtered spent media using Liquid Chromatography-Mass Spectrometry (LC-MS).

For GC-MS, 10 μL of 1.0 mg/mL ribitol and 10 μL of 1.0 mg/mL nonadecanoic acid in hexane were added to 100 mg cell samples. Then, 0.35 mL of ice-cold methanol was added to the samples, vortexed for 3 minutes and placed into an ice-cold sonication water bath for 10 min at full output power. Two 2.4 mm metal beads were added to the samples and the biomass was homogenized using a Bead Mill 24 Homogenizer (Thermo Fisher Scientific, Inc., Waltham, MA, USA). Next, 0.35 mL of chloroform was added, and the samples were vortexed for 3 min followed by the addition of 0.30 mL of water and another 3 minutes of vortex before the samples again were placed into an ice-cold sonication water bath for 10 min at full output power. The samples were vortexed for 3 min and then centrifuged for 7 min at maximum speed (16,000 g). Three hundred microliters of upper layer (polar fraction) and 200 μL of lower layer (nonpolar fraction) were transferred to GC-MS vials. Samples were dried in a speed-vac concentrator for 10 h and then derivatized following established protocols [15]. The derivatized samples were analyzed at the Iowa State University W.M. Keck Metabolomics Research Laboratory (RRID:SCR_017911). GC-MS analyses were performed with an Agilent 6890 gas chromatograph coupled to a model 5973 Mass Selective Detector (Agilent Technologies, Santa Clara, CA). The column used was HP-5MS 5% Phenyl Methyl Silox with 30 m × 250µM × 0.25 µm film thickness (Agilent Technologies). The mass data were collected in the range from m/z 40 to m/z 800. Chromatograms were analyzed using NIST AMDIS (Automated Mass Spectral Deconvolution and Identification System) software.

For LC-MS analysis, 100 mL aliquots of filtered spent media were mixed 1:1 v/v with ethyl acetate, mixed, allowed to separate and the organic phase was collected using a separatory funnel. The organic phase was evaporated using a rotavap, the recovered extract was resuspended in ethanol to a final concentration of 2 mg of extract / mL of ethanol and the internal standard 1,2-didecanoyl-sn-glycero-3-phosphocholine was added to each sample at a final concentration of 1 µM. The samples were analyzed in positive and negative-ion mode at the W.M. Keck Metabolomics Research Laboratory with an Agilent Technologies 1290 Infinity Binary Pump UHPLC instrument equipped with an Agilent Technologies Eclipse C18 1.8 μm x 2.1 mm × 100 mm analytical column coupled to an Agilent Technologies 6540 UHD Accurate-Mass Q-TOF mass spectrometer.

### Statistical Methods

Cultures were monitored using OD_600_ measurements performed in triplicates and normalized against negative controls or blanks. In comparative expression experiments Log2 Fold Changes were calculated using the following formula:

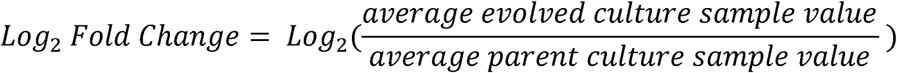

Mean comparisons for two samples were conducted using the Student’s t-test. Mean comparisons of multiple samples were completed with one-way ANOVA and *p*-values obtained from the all-pair Tukey Kramer test in JMP Pro 16. Propagation of error formulas were used to estimate uncertainties [22].

## Results and Discussion

### TOD Characterization

Thermal oxo-degradation of HDPE, the characterization of its products, and the ability of the substrates to support microbial growth were discussed in a previous publication [5]. This previous work characterized TOD product that was collected in three stage fractions with condensers and their ability to support microbial growth. The molecules in the 200-600 Da range were found to support the best growth. Therefore, we modified the TOD reactor to produce a single fraction with a molecular weight distribution close to the 200-600 Da range and avoid condensers to reduce the energy demand. The single product from this condition was collected as a waxy single stage fraction at room temperature. The TOD_HDPE wax was characterized again via FTIR (Figure 2A-B), acid titration (Figure 2C), GPC (Figure 2D), and elemental analysis (Figure 2E). The FTIR analysis is consistent with the prior report of the TOD fractions, with the characteristic peak for carbonyl groups at 1650-1850 cm ¹ demonstrating the introduction of oxygenated functionalities such as carboxylic acids and aldehydes. The carbonyl index (CI) of 0.9 ± 0.1 (Figure 2B) is indistinguishable from the CI of the individual wax or liquid fractions. However, the total acid number (TAN) of 0.93 ± 0.08 (Figure 2C) is decreased relative to the prior configuration. The GPC chromatogram (Figure 2D) indicates a molecular weight distribution with >90% of the molecules ranging between 268-988 Da, corresponding to 19 to 68-carbon molecules, with the major peak at around 490 Da. In the prior configuration, the two waxy fractions both had major peaks around 600 Da and the liquid fraction’s major peak was just over 200 Da.

**Figure 2.**
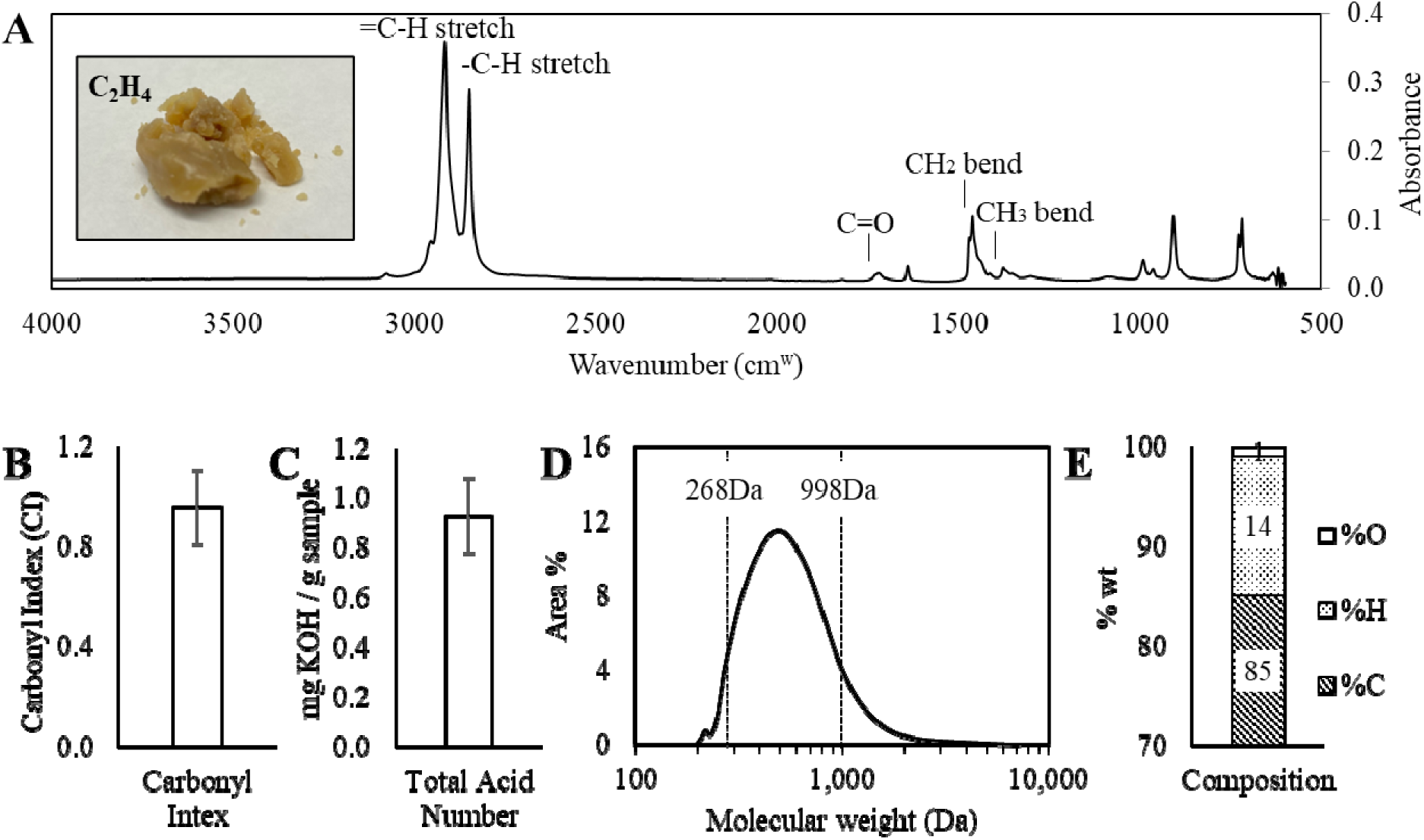
Characterization of TOD_HDPE product. (A) FTIR spectra shows the presence of hydrocarbons with terminal alkenes (=C-H) and oxygenated functionalities with carbonyl groups (C=O). (B-C) The carbonyl index (CI) and total acid number (TAN) support the oxidation of HDPE cracking products during thermal deconstruction and are consistent with previous reports. Error bars represent standard deviation of measurements. (D) GPC shows a molecular weight distribution with 90% of the molecules falling within the 268-988 Da range, corresponding to 19 to 68-carbon molecules. (E) The elemental composition corresponds to a C_23_H_44_O_1_ stoichiometric ratio.

The elemental composition (Figure 2E) indicates an oxygen content of 0.7 ± 0.02 wt.% decreased relative to the 2.1±0.5 and 2.5±0.9 observed for the two waxy fractions. The overall characterization confirms the cracking and oxidative effects of the thermal oxo-degradation of HDPE, while highlighting a slight decrease in oxygenation compared to the products from the previously published TOD products due to changes in reactor configuration and operating conditions.

### Specific growth rate in TOD_HDPE improved by over 100% during evolution

Wild-type *C. maltosa* could take over 30 hours to reach an OD_600_ of 0.6 in TOD_HDPE from the new TOD reactor configuration collected in a single fraction. After 14 serial transfers in media containing TOD_HDPE, the evolved *C. maltosa* population was able to obtain an OD_600_ of 0.6 in approximately 9 hours while maintaining the starting OD600 constant. Twenty distinct isolates from the evolved population were studied in four sets of mixed cultures, each consisting of five strains to maximize the number of isolates we could test with the available material. The specific growth rates of the co-cultures ranged from 0.022 to 0.069 h^-1^. The five strains composing the group with the fastest growth rate were tested individually and the fastest growing strain had a specific growth rate of 0.098h^-1^, an increase of over 100% relative to the parent strain (Table 1). This strain was selected as the final evolved strain and named *C. maltosa* ERO1.

**Table 1.** The evolved strain characterized here (ERO1) was chosen out of 20 isolated colonies from the evolved population. ERO1 shows a greater than 100% increase (p-value 0.03) in specific growth rate in TOD_HDPE relative to its parent.

| Isolates | Specific Growth Rate (h<sup>-1</sup>) |
| --- | --- |
| *Parent Strain* |  |
| NRLL Y-17677 | 0.029 ± 0.004 |
| *Evolved Population: Co-cultured Colonies* |  |
| 1-5 | 0.041 ± 0.008 |
| 6-10 | 0.06 ± 0.02 |
| 11-15 | 0.02 ± 0.04 |
| 16-20 | 0.07 ± 0.01 |
| *Evolved Population: Single Colonies* |  |
| 16 (ERO1) | 0.098 ± 0.034 |
| 17 | 0.04 ± 0.05 |
| 18 | 0.059 ± 0.009 |
| 19 | 0.010 ± 0.005 |
| 20 | 0.007 ± 0.012 |

### The evolved strain showed off-target benefits with minimal trade-offs

While the directed evolution was intended to select for increased growth rate of *C. maltosa* in TOD_HDPE, the evolved strain also showed improvements in conditions not directly selected for during adaptive evolution. For example, its growth was also studied in the same set of model compounds used in a previously published yeast survey that characterized the substrate range of oleaginous and nonconventional yeast species [7], and from which *C. maltosa* was chosen for TOD_HDPE utilization. The model compounds included C14, C18 and C22 carboxylic acids, alcohols, alkane, alkenes and esters. Compared to the parent strain, the evolved strain showed significant increase in total growth in 48-hour cultures in glucose and esters at the expense of reduced total growth in the C12 alkene and C18 alkane (Figure 3A). The evolved strain also outperformed the parent strain in growth rate and total growth when grown in the TOD products from a 1:1 mixture of HDPE/Polypropylene feedstock (TOD_HDPE/PP), as shown in Figure 3B. The preliminary composition of TOD_HDPE/PP can be found in the Supplemental Table S1. Finally, the evolved strain also outperformed the parent strain in culture conditions where the traditional nitrogen source (AmSO_4_) had been replaced with human urine (Figure 3C), which shows promise for broader applications with wastewater feedstocks.

**Figure 3.**
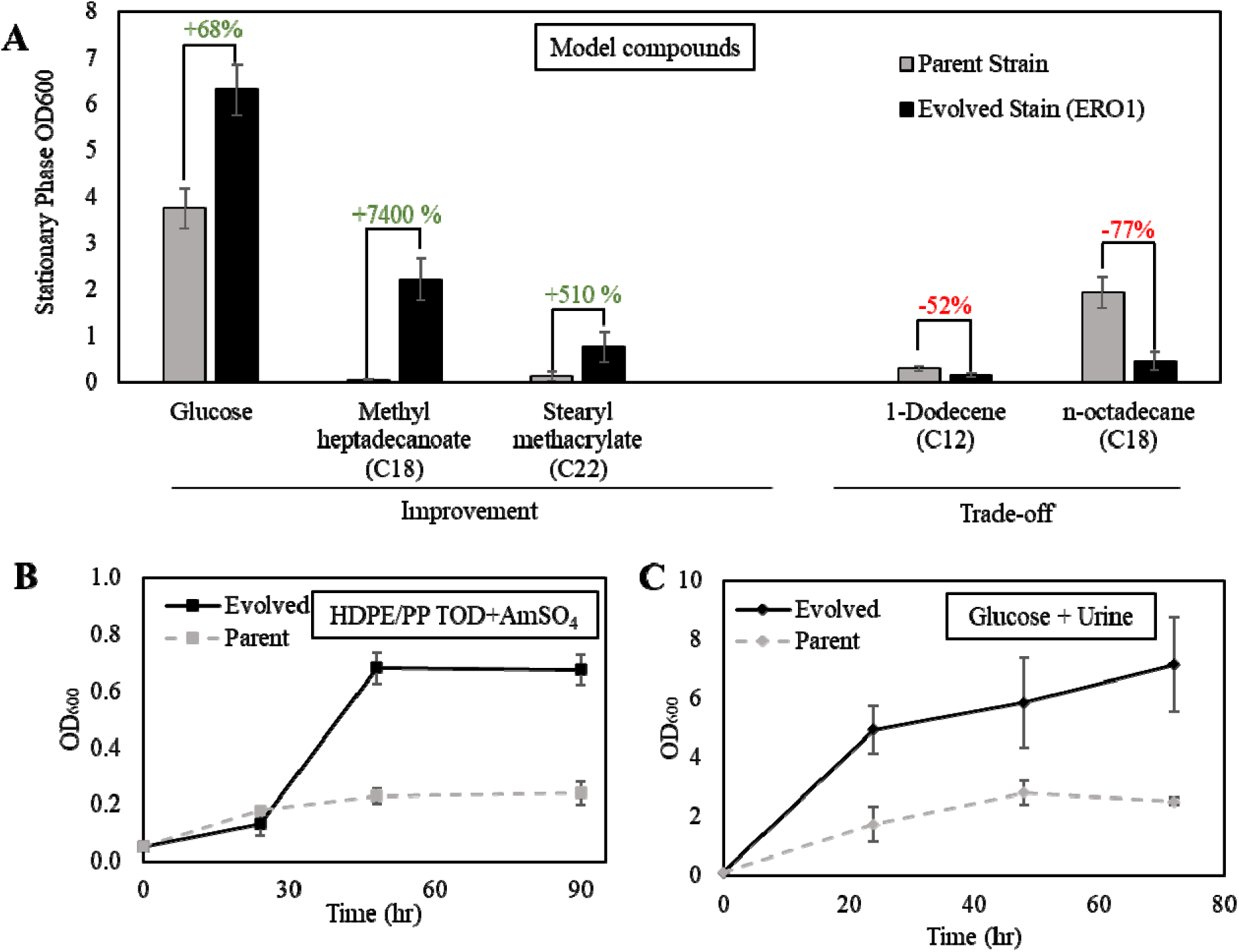
(A) Stationary phase OD_600_ of parent and evolved strains grown in 0.05M model compounds where the differences between strains were significant (p-value < 0.05) shows a trade-off between hydrocarbons and esters. (B) Flask growth curves with 0.5% w/v TOD_HDPE/PP (composition in Table S1) as the sole carbon sources. The evolved strain shows significant improvement in TOD_HDPE/PP utilization. (C) Growth curve comparison between parent and evolved strain in 2% w/v glucose with 15% v/v urine as a nitrogen source shows improved growth in the evolved strain.

Understanding these changes in growth profile may also help future design strategies for the development of an industrial strain or more comprehensive bioprocess. The trade-off between growth on hydrocarbons and molecules with oxygenated functionality could shed some light into the underlaying metabolic pathways involved in the utilization of TOD_HDPE, which is a complex mixture. The improved growth in TOD_HDPE/PP and urine conditions are valuable because of their implications for future development of the plastic upcycling process to process additional types of plastics and leverage other waste streams as nutrient sources [23].

### *C. maltosa* solubilization of hydrophobic molecules improved in evolved strain

TOD_HDPE is a solid and hydrophobic feedstock that floats on the surface of the aqueous media during cultures (Figure S1) and *C. maltosa* can use it as its sole carbon source. Not only can *C. maltosa* grow on TOD_HDPE, but our previous work also showed that the parent strain can grow on TOD substrates as well as it does with glucose [5]. Although it was expected that mass transfer limitations would occur in this system, results demonstrate that *C. maltosa* has native mechanisms to overcome such limitations and make the carbon source available in the aqueous phase.

To test the solubilization activity of *C. maltosa*, the solubility of four model compounds was measured in glucose media without cells and in filter-sterilized spent glucose media from parent and evolved cultures, all incubated for 48 hours (Figure 4). The four model compounds (1-tetradecanol, 1-octadecanol, n-tetradecane and n-octadecane) were confirmed to have very low solubility in the growth media. Specifically, each compound was added to the no cell control media individually, vortexed and filtered and their abundance assessed by GC-MS; all four were below the detection limit. This process was repeated using filtered spent media obtained after 48 hours of utilization by the parent and evolved strains separately. While the 1-tetradecanol and n-tetradecane remained below the detection limit, the baseline solubilization activity of the parent strain is evidenced by the increase of 1-octadecanol and n-octadecane solubility to roughly 13 mM. The solubilization activity of the evolved strain is increased such that 1-tetradecanol became detectable in the aqueous phase and the solubility of 1-octadecanol was increased relative to the parent strain.

**Figure 4.**
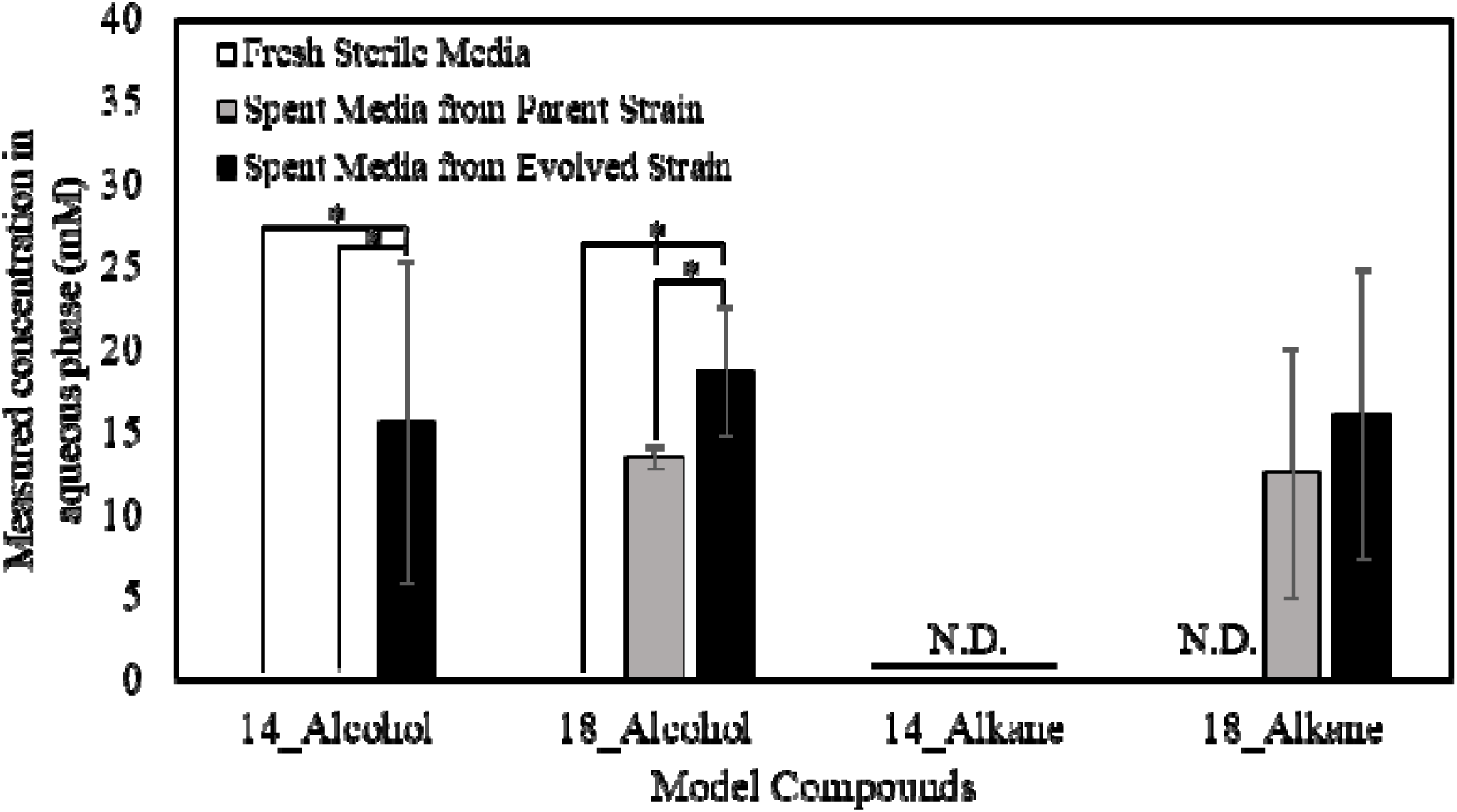
Solubilities of model compounds representative of TOD_HDPE composition show that otherwise insoluble molecules, which are not detected in solution in sterile media, go into solution in spent media from the parent and evolved strains of *C. maltosa* cultured in YNB + 2% glucose for 48 hours. Solubilities were tested in YNB media at room temperature. N.D. = Not Detected and asterisks indicate significant differences (p-value < 0.05).

The results in Figure 4 show that biological products of *C. maltosa* solubilize otherwise insoluble molecules, increasing their bioavailability in the aqueous media. The fact that this solubilization activity increased in the evolved strain suggests that this is a mechanism to overcome mass transfer limitations. We hypothesized that the mechanism behind the solubilization activity involved the secretion of biosurfactants because surfactants are known solubilize hydrophobic molecules [24].

### *C. maltosa* biosurfactant properties are altered in evolved strain

The results described above used spent media obtained during growth on glucose. Using a similar procedure, spent media was collected during growth on TOD_HDPE in order to characterize the proposedTOD_HDPE biosurfactants.

Biosurfactants are known to reduce the surface tension of liquids and emulsify non-polar solvents [25] and thus the surface tension and emulsification index were measured (Figure 5). Consistent with the increased solubilization of model compounds, the spent media from both the parent strain and evolved strain grown on glucose showed a significant decrease in surface tension relative to the media control (Figure 5A). Similar effects were observed using TOD_HDPE as carbon source instead of glucose.

**Figure 5.**
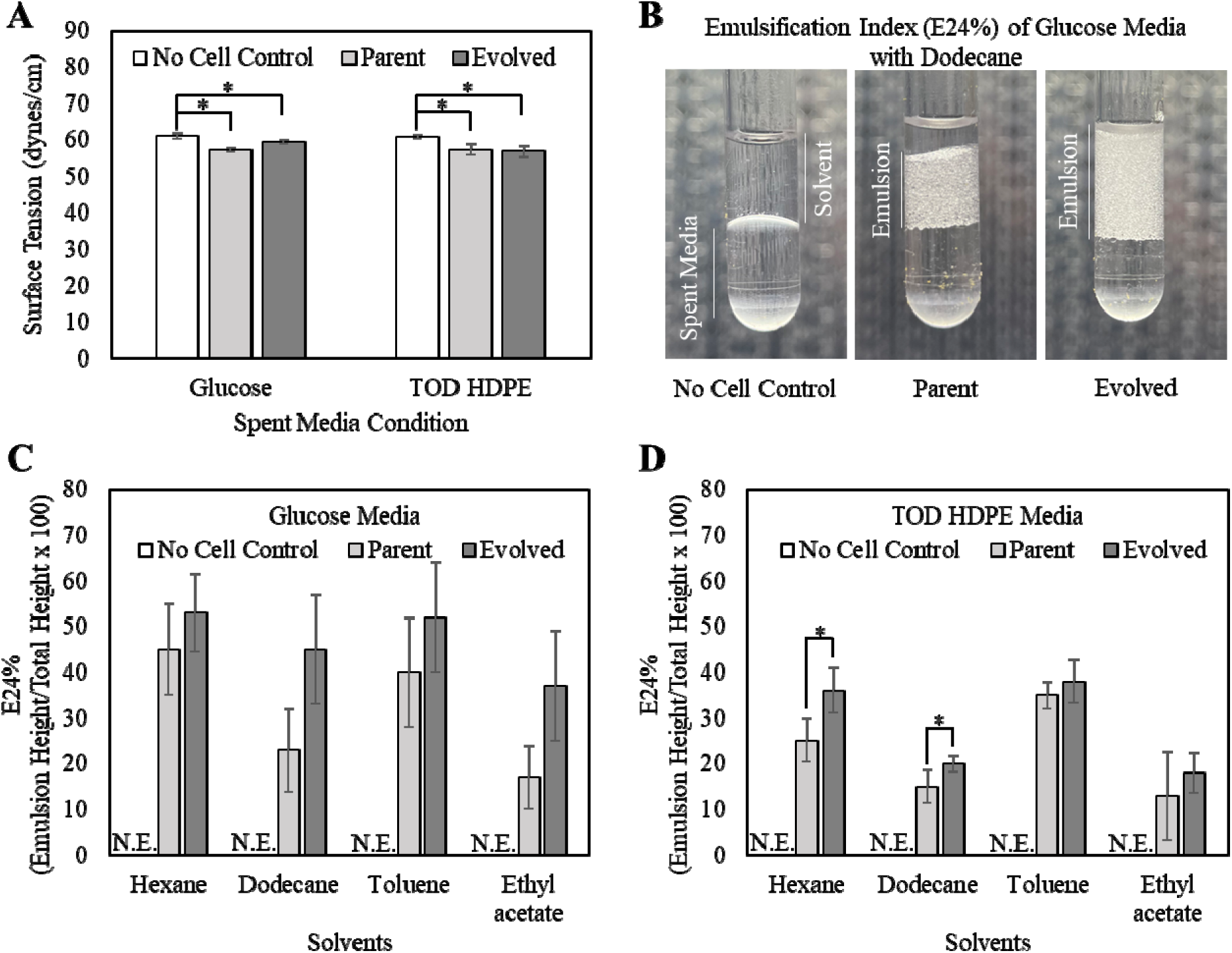
Decreases in surface tension (A) relative to no cell controls and the formation of emulsions (B-D) indicate the presence of biosurfactants in both parent and evolved cultures grown with glucose and TOD_HDPE. (B) The Emulsification Index (E24%) is the height ratio of emulsion relative to the total volume in a 1:1 filtered spent media:solvent mixture. This figure shows one replicate of the dodecanol condition presented in Figure 5C. (C) E24% measurements for glucose media conditions show no difference between parent and evolved. (D) E24% measurements for TOD_HDPE media conditions show higher emulsification of alkane by the evolved strain’s spent media. N.E. = No Emulsion and asterisks indicate significant differences (p-value < 0.05).

The biosurfactant presence was further characterized by measurement of the emulsification index (E24) (Figure 5B). While mixing of the media-only control and dodecanol results in two stable phases, the same process with glucose spent media yields an emulsified layer at the interface of the aqueous and organic layers. The height of this emulsified layer is indicative of the biosurfactant activity. Consistent with the measurements of molecule solubilization and surface tension, the emulsification indeces of the glucose spent media (Figure 5C) and TOD HDPE spent media (Figure 5D) of both the parent and evolved strains were increased relative to the media only control across all four solvents (hexane, dodecane, toluene and ethyl acetate).

In addition to detecting the presence of biosurfactants, the results also indicate that the adaptative laboratory evolution affected the properties of the biosurfactants. When grown in TOD_HDPE, the evolved strain showed significantly increased emulsification activity for the two alkane solvents tested (hexane and dodecane) relative to the parent strain. The fact that the evolution process resulted in better emulsification of alkanes and not toluene or ethyl acetate is relevant because alkanes are one of the principal components of TOD_HDPE. This observation could indicate a change in relative abundances of the biosurfactants to favor alkane utilization, a change in the amount of biosurfactants secreted or both.

The presence of biosurfactants was detected indirectly though their effect on the properties of spent media. To identify the potential classes of biosurfactant present, the modified Lowry, saponification, anthrone and rhamnose tests were performed in the TOD_HDPE spent media to identify the presence proteins, lipids, carbohydrate and rhamnose [26], [27]. The results, summarized in Table 2, show the presence of proteins, lipids and carbohydrates, and the absence of rhamnose, in both parent and evolved cultures. Table 2 also shows the absence of all components, except lipids, in the no cell control. The presence of lipids in the no cell control, originating from the TOD_HDPE substrate, obscures the results in the culture conditions. However, oleaginous yeasts like *C. maltosa* are known to produce lipids, and therefore it is likely that there are additional lipids in the spent media from the cultures [28]. Based on these results, the biosurfactants produced by *C. maltosa* could be glycolipids, lipopeptides and or proteins, but not rhamnolipids [29]. Special attention should be given to the protein and lipopeptide candidates because the evolved strain showed significantly higher concentration of protein in the spent media relative to the parent strain.

**Table 2.** The biochemical tests performed on the spent media showed the presence of proteins, lipids, and carbohydrates in parent and evolved cultures.

| Test | No cell control | *C. maltosa*<br>Parent | *C. maltosa*<br>Evolved |
| --- | --- | --- | --- |
| Proteins ($\mu\text{g/mL}$) – Modified Lowry | Negative | $78 \pm 7$ | $110 \pm 20$ |
| Lipids – Saponification Test | Positive | Positive | Positive |
| Carbohydrates ($\text{OD}_{620}$) – Anthrone | Negative | $0.11 \pm 0.03$ | $0.13 \pm 0.01$ |
| Rhamnose Test | Negative | Negative | Negative |

### Proteome analysis points to cellular adaptations confirmed by electron microscopy

The biochemical tests showed significantly higher concentration of proteins in the spent media from the evolved strain relative to the parent strain. It was hypothesized that the changes in biosurfactants that led to improved emulsification of alkanes could be an increase in the secreted lipopeptides or proteins acting as biosurfactants. A shotgun proteomic analysis was performed on the parent and evolved cells harvested during mid-exponential phase and on the spent media from stationary phase in TOD_HDPE to identify the potential biosurfactants with differing abundance in the parent and evolved strains. The analysis did not identify any differentially expressed proteins with known biosurfactant activity, but it did identify two proteins in the spent media and 18 intracellularly expressed proteins with significant difference in abundance between parent and evolved strains (Table 3). Complete list of identified proteins can be found in Table S2.

**Table 3.**
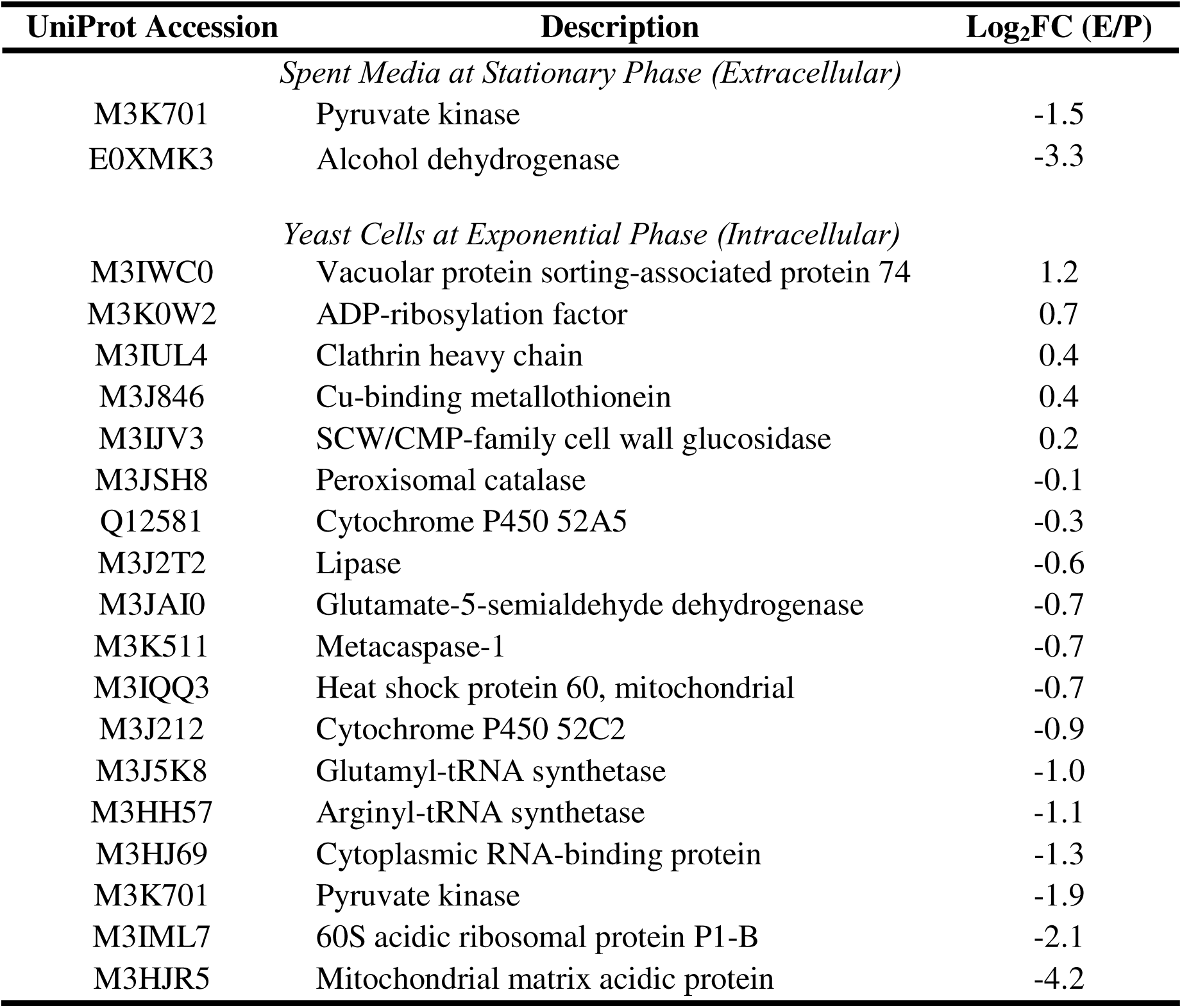
Identified proteins with significant difference in abundance between parent and evolved strains provide insights into cellular and metabolic functions involved in TOD_HDPE utilization. The Fold Change (Log_2_FC) shows the difference in evolved strain relative to parent.

In the spent media, pyruvate kinase and alcohol dehydrogenase were less abundant in the evolved conditions. The private kinase was also less abundant in the cell extracts from the evolved strain. These enzymes play crucial roles in sugar metabolism and the absence of carbohydrates in the carbon source could contribute to the reduced presence of these enzymes [30], [31].

Five intracellular proteins showed higher abundance in the evolved strain, with three findings discussed here. The vacuolar protein sorting-associated protein 74 is involved in the sorting of catabolic enzymes, which may play a role in the metabolism of the complex mixture that is TOD_HDPE [32]. The clathrin heavy chain protein forms part of the protein system that initiates intracellular vesicle formation, which facilitate transport between membrane-bound compartments like vacuoles and peroxisomes, also relevant to TOD_HDPE metabolism [33], [34], [35]. Lastly, the SCW/CMP-family cell wall glucosidase plays a role in the formation and changes to polysaccharides in the cell wall, which are known to change in *C. maltosa* in the presence of alkanes as part of the process to form canals that may facilitate transport and metabolism of hydrophobic substrates [36], [37]. Cross-sectional TEM images of *C. maltosa* show potential confirmation of the cellular mechanisms suggested by the proteomic analysis are induced by TOD_HDPE. Figure 6 shows confirmation of canal formation when grown in the presence of TOD_HDPE and Figure S2 show the abundance of compartments induced by TOD_HDPE, including peroxisomes, vacuoles and vesicles.

**Figure 6.**
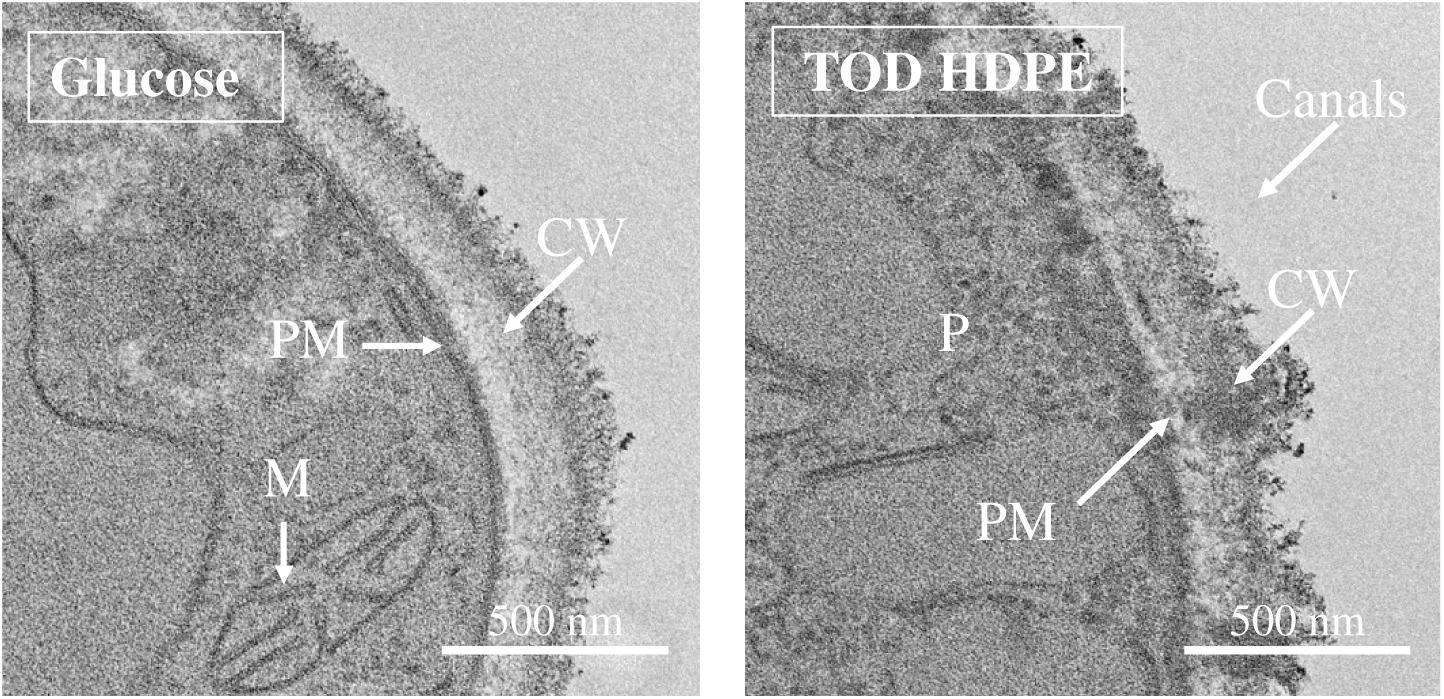
TEM cross-sectional images of the *C. maltosa* parent strain grown in glucose and in TOD_HDPE show the formation of cell wall canals and peroxisomes and a reduction in cell wall thickness in the presence of TOD_HDPE. Similar structures were observed in parent and evolved strains (not shown). PM = plasma membrane, CW = cell wall, M = mitochondria.

Thirteen proteins were less abundant in the evolved strain relative to the parent strain. Surprisingly, two Cytochrome P450 proteins, which play a role in alkane metabolism, were among the proteins with reduced abundance [38]. It was also surprising that a peroxisomal catalase and a lipase, both involved in lipid metabolism, were also less abundant in the evolved strain extracts [39], [40].

### Membrane permeability modulated by ergosterol may play a role in bioconversion

Overcoming the mass transfer limitations posed by the hydrophobic substrate is only the first challenge in the uptake of TOD_HDPE. The molecular weight distribution of the TOD_HDPE showed that most molecules in the substrate are >19 carbons long. Transporting molecules that big inside the cells is usually problematic [41], [42]. Still, *C. maltosa* can metabolize those molecules[41], [42]. To better understand the transport of these molecules into the cells, the membrane permeability was studied. Membrane permeability is an important metric to study because changes in fluxes across the membrane could substantially impact important metabolic processes of the cell.

SYTOX green was used to measure permeability because it exhibits a >500-fold fluorescence enhancement upon binding to nucleic acids and it must pass through the membranes to do so [43]. Therefore, the fluoresce intensity of a normalized group of cells can be taken as a measure of how much dye was able to access the nucleic acids. Using these principles, our assay compared the permeability of the two *C. maltosa* strains grown in glucose before and after exposure to TOD_HDPE and a known membrane permeabilization agent (Triton X-100) for one hour (Figure 7A). There was no significant difference in permeability between the parent and evolved strain and both strains showed significant increase in permeability when exposed to Triton X-100. However, only the evolved strain showed increased permeability when exposed to TOD_HDPE. The membrane permeability of parent and evolved cells grown TOD_HDPE cultures for 48 hours was also measured (Figure 7B). The results were consistent with those of the cells grown in glucose and exposed to TOD_HDPE for one hour.

**Figure 7.**
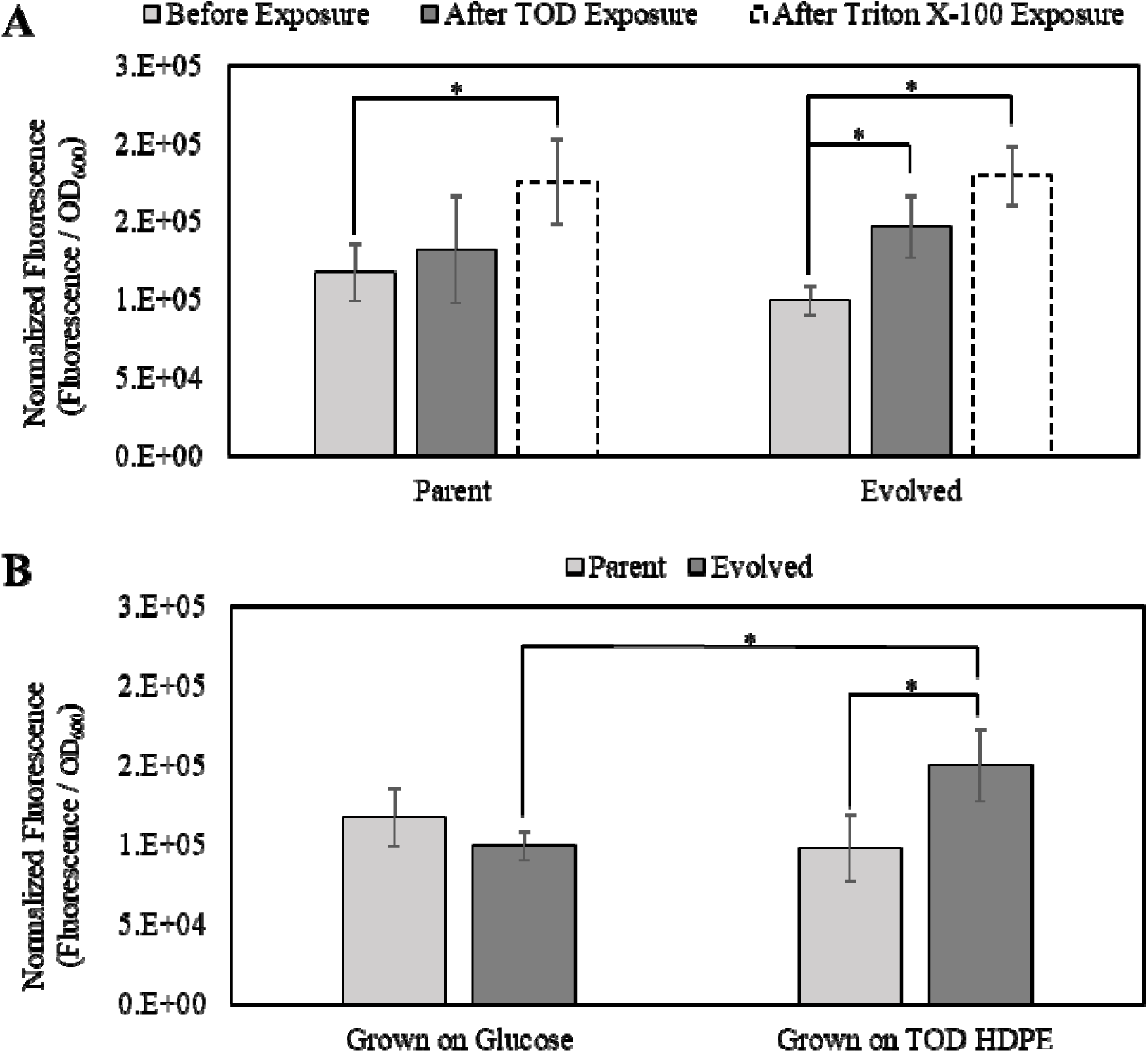
(A) The permeability of the evolved strain’s membrane increases in response to TOD_HDPE exposure for one hour, while the parent strain’s membrane does not. (B) The permeability of the evolved strain after 48-hour growth in TOD_HDPE is consistent with the permeability after only one hour, while the parent strain remains unchanged (p-value < 0.05).

Increase in membrane permeability is usually interpreted as a sign of membrane damage [44], [45]. However, in this case, there is no difference in permeability between the parent strain grown in TOD_HDPE relative to glucose, ruling out that TOD_HDPE causes membrane damage. Only the evolved strain increases its permeability in response to TOD_HDPE, and the magnitude of the permeability increase is comparable after one hour or 48 hours of exposure to TOD_HDPE, instead of increasing with time and leading to cell death. These facts suggest that the increased permeability may contribute to the improved phenotype as a beneficial cellular adaptation. As Orgel’s second rule states, evolution is cleverer than we are [46].

The mechanism for the change in membrane permeability may be explained by the metabolite analysis of the cells. A non-targeted metabolite analysis of the yeast cells from the parent and evolved strains grown in TOD_HDPE was conducted with GC-MS. The results, shown in Table 4, identified six molecules with a significant difference (p-value <0.05) in abundance between strains. The molecule with the largest decrease in the evolved strain was ergosterol (Log_2_FC -2.06). Ergosterol is the most abundant sterol in fungal cell membranes, and it is known to regulate their permeability and fluidity [47], [48].

**Table 4.** Non-targeted GC-MS metabolite analysis of the yeast cells at stationary phase identified six molecules with significant differences in abundance between parent and evolved strains of C. maltosa.

| Name | CAS No. | Function | Formula | Log<sub>2</sub>FC (E/P) |
| --- | --- | --- | --- | --- |
| Ergosterol | 57-87-4 | Membrane stability | C<sub>28</sub>H<sub>44</sub>O | -2.06 |
| 1-Monooleoylglycerol | 111-03-5 | Surfactant | C<sub>21</sub>H<sub>40</sub>O<sub>4</sub> | -1.91 |
| D-Glucitol | 50-70-4 | Osmoregulation | C<sub>6</sub>H<sub>14</sub>O<sub>6</sub> | -1.50 |
| Glycerol | 56-81-5 | Lipid backbone | C<sub>3</sub>H<sub>8</sub>O<sub>3</sub> | -1.15 |
| 4-Hydroxybutanoic acid | 591-81-1 | Intermediate | C<sub>4</sub>H<sub>8</sub>O<sub>3</sub> | -0.66 |
| 1-Hexatriacontene | 61868-14-2 | Alkene | C<sub>36</sub>H<sub>72</sub> | 1.82 |

### The laboratory evolution affected the metabolites secreted by *C. maltosa*

As a non-conventional yeast, the metabolite profile of *C. maltosa* is mostly unknown. To study the molecules secreted during growth in TOD_HDPE and how they were affected by the laboratory evolution, the spent media of the parent and evolved strains was subjected to LC-MS in positive and negative mode. Given the complex composition of TOD_HDPE, the media of a no cell control incubated alongside the cultures was also studied. Table 5 lists the molecules absent in the TOD_HDPE no cell control but present in the parent and/or evolved cultures, and the log fold change between strains for molecules present in both cultures with significant difference in abundance.

**Table 5.**
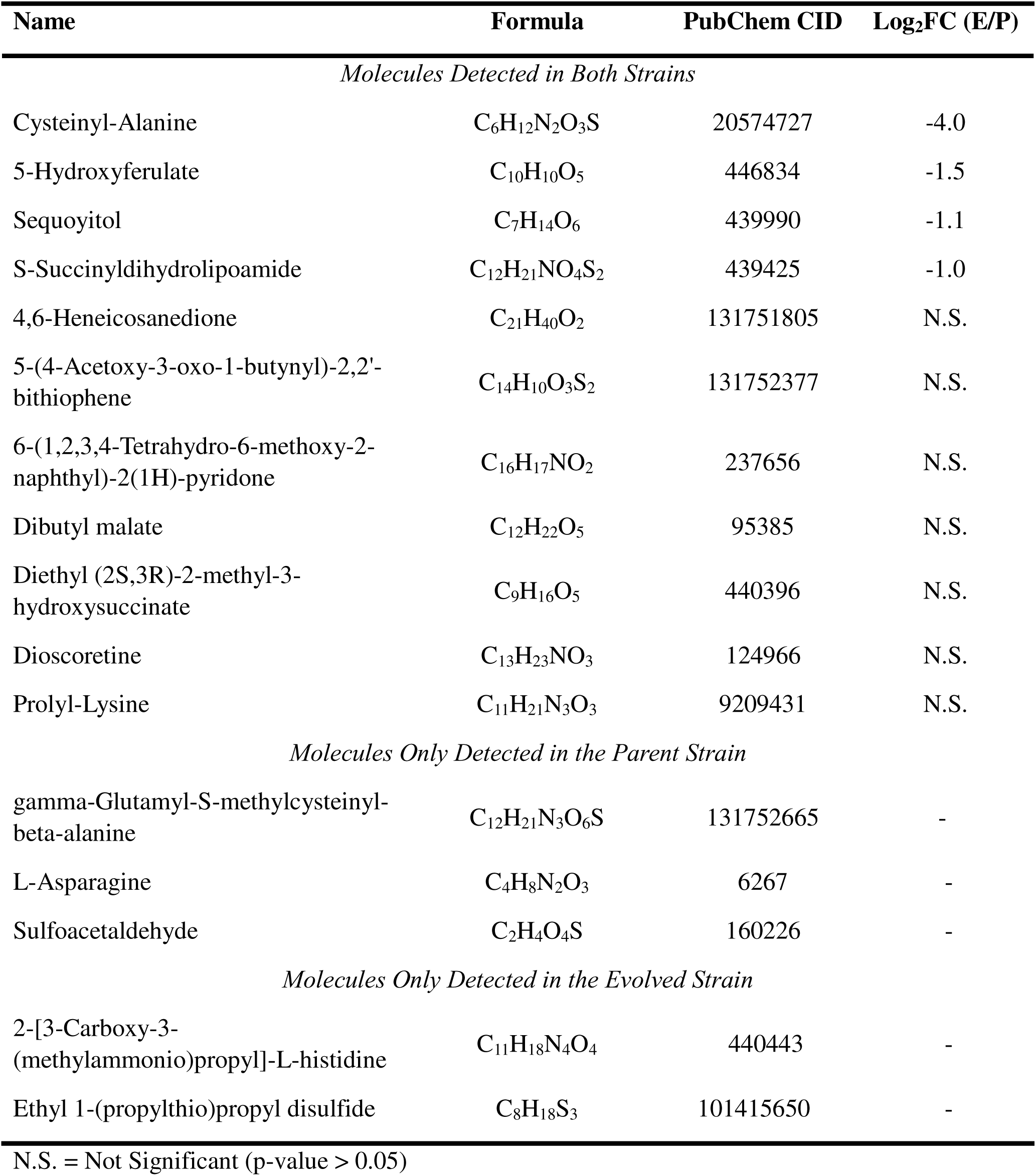
Molecules of biological origin identified via LC-MS present only in the spent media from parent and/or evolved cultures of *C. maltosa* grown in TOD_HDPE and absent in no cell controls.

Nine of the identified molecules in the parent culture were either significantly less abundant in the evolved culture or present in the culture of only one of the strains. The molecules with reduced abundance were S-succinyldihydrolipoamide, sequoyitol, 5-hydroxyferulate and cysteinyl-alanine. S-succinyldihydrolipoamide is a fatty acid derivative involved in membrane stabilization, sequoyitol is a methylated inositol with possible role in phospholipid synthesis regulation, and Cysteinyl-Alanine is an incomplete breakdown product of protein catabolism [49], [50], [51], [52]. The 5-hydroxyferulate is a suspicious finding because it is an intermediate in the conversion of ferulic acid to sinapic acid in lignin biosynthesis [53]. Interestingly, however, the enzyme that produces 5-hydroxyferulate is dependent on cytochrome P450, and our proteomic analysis showed significant reduction in cytochrome P450 proteins in the evolved strain.

## Conclusions

*C. maltosa* is a poorly studied organism with remarkable mechanisms to utilize industrially and environmentally relevant hydrophobic compounds and an untapped potential for biotechnology. One of its most interesting qualities is the ability to force insoluble molecules into solution to make them bioavailable in aqueous media. Due to its solubilization activity and its ability to metabolize long-chain alkanes, fatty acids and fatty alcohols, *C. maltosa* is a promising microbial cell factory for plastic upcycling via bioconversion of depolymerized plastic wastes. Adaptive laboratory evolution made *C. maltosa* a better microbial platform for bioconversion by increasing the specific growth rate in polyethylene-derived feedstock by over 100%. A cellular and biochemical comparison of the parent and evolved strains revealed that the improved phenotype may be attributed to improved solubilization activity due to changes in secreted biosurfactants and an increase in membrane permeability due to a decrease in ergosterol in the presence of the depolymerized polyethylene. The underlying mechanism that contributed to the improved phenotype characterized in this study may inform rational engineering strategies to develop industrial strains for waste valorization and treatments or provide insights for bioremediation applications.

## Acknowledgements

We acknowledge the Iowa State University W.M. Keck Metabolomics Research Laboratory (RRID:SCR_017911), Protein Facility, and Roy J. Carver High Resolution Microscopy Facility for providing analytical, proteomic, and microscopy instrumentation and guidance. We thank Dr. Lucas J Showman, Dr. Matthew W Breitzman, Joel Nott and Tracey Stewart for their assistance and support.

