## Supplemental Information for "Adaptive evolution of *Candida maltosa* improves the bioconversion of depolymerized plastic feedstock by targeting biosurfactant production"

**Supplemental Materials**

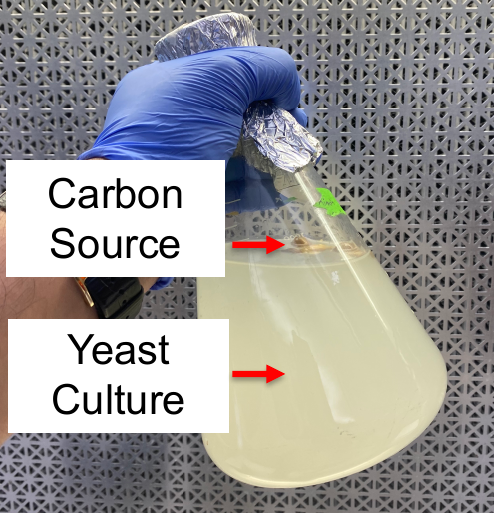

**Figure S1.** TOD_HDPE shown floating at the top of the aqueous phase post culture.

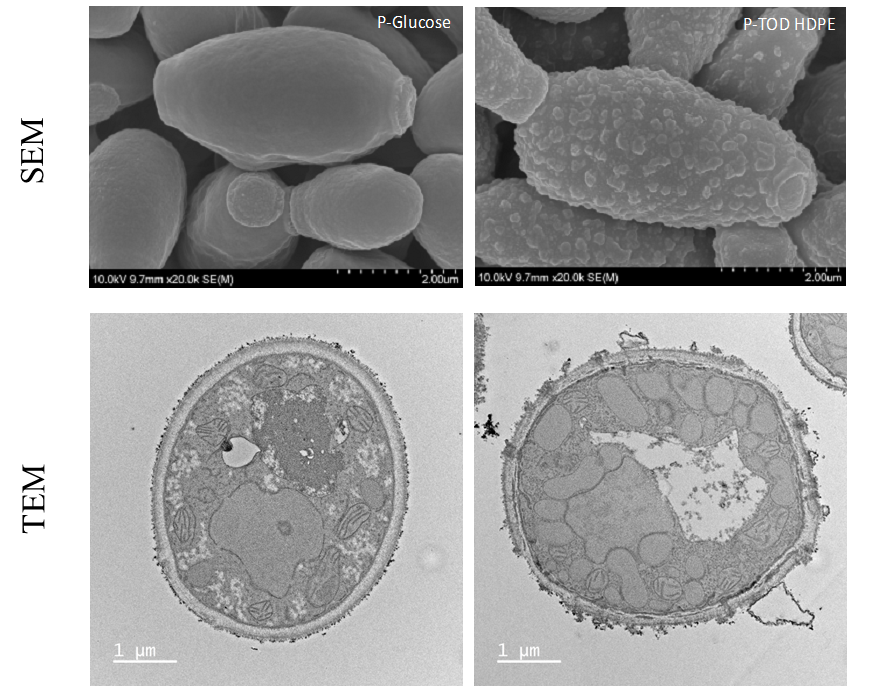

**Figure S2.** SEM and cross-sectional TEM micrographs of *Candida maltosa* show proliferation canals and intracellular compartments, including vacuoles and peroxisomes, induced by the presence of TOD_HDPE.

**Table S1.** TOD_HDPE/PP composition. Identification and quantification of the largest 30 peaks on GC-MS/FID measured of the GC-detectable fraction; the products are a mixture of alcohols, aldehydes, alkanes, alkenes, and alkadienes.

| **Compound** | **Concentration (wt.%)** |
| --- | --- |
| *Alcohol* | |
| 2-nonen-1-ol | 0.32 |
| 1-octanol, 2-butyl | 1.04 |
| 2-tridecen-1-ol | 0.40 |
| 18-nonadecen-1-ol | 0.58 |
| 1-heneicosanol | 1.38 |
| behenic alcohol | 1.52 |
| 1-heneicosanol | 1.55 |
| n-tetracosanol-1 | 0.91 |
| Octacosanol | 1.58 |
| *Aldehyde* | |
| 10-undecen-1-al, 2-methyl- | 0.35 |
| 10-undecenal | 0.34 |
| *Alkane* | |
| dodecane | 0.91 |
| hexadecane | 0.69 |
| 13-oxabicyclotridecane | 0.46 |
| 2-methyltetracosane | 0.64 |
| octadecane | 0.49 |
| eicosane | 0.54 |
| heptacosane | 0.73 |
| *Alkene* |  |
| 1-dodecene | 1.64 |
| 1-tridecene | 3.26 |
| 1-tetradecene | 2.95 |
| 1-pentadecene | 2.43 |
| cetene | 1.81 |
| 1-heptadecene | 1.50 |
| 1-nonadecene | 1.21 |
| 5-eicosene | 1.23 |
| *Alkadiene* | |
| 1,11-dodecadiene | 2.72 |
| 1,12-tridecadiene | 1.34 |
| 1,15-hexadecadiene | 0.57 |
| 1,19-eicosadiene | 0.56 |
